# ZARP: An automated workflow for processing of RNA-seq data

**DOI:** 10.1101/2021.11.18.469017

**Authors:** Maria Katsantoni, Foivos Gypas, Christina J. Herrmann, Dominik Burri, Maciej Bak, Paula Iborra, Krish Agarwal, Meric Ataman, Anastasiya Börsch, Mihaela Zavolan, Alexander Kanitz

## Abstract

RNA sequencing (RNA-seq) is a crucial technique for many scientific studies and multiple models, and software packages have been developed for the processing and analysis of such data. Given the plethora of available tools, choosing the most appropriate ones is a time-consuming process that requires an in-depth understanding of the data, as well as of the principles and parameters of each tool. In addition, packages designed for individual tasks are developed in different programming languages and have dependencies of various degrees of complexity, which renders their installation and execution challenging for users with limited computational expertise. The use of workflow languages and execution engines with support for virtualization and encapsulation options such as containers and Conda environments facilitates these tasks considerably. Computational workflows defined in those languages can be reliably shared with the scientific community, enhancing reusability, while improving reproducibility of results by making individual analysis steps more transparent.

Here we present ZARP, a general purpose RNA-seq analysis workflow which builds on state-of-the-art software in the field to facilitate the analysis of RNA-seq data sets. ZARP is developed in the Snakemake workflow language using best software development practices. It can run locally or in a cluster environment, generating extensive reports not only of the data but also of the options utilized. It is built using modern technologies with the ultimate goal to reduce the hands-on time for bioinformaticians and non-expert users. ZARP is available under a permissive Open Source license and open to contributions by the scientific community.

## Introduction

Recent years have seen an exponential growth in bioinformatics tools [1], a large proportion of which are dedicated to High Throughput Sequencing (HTS) data analysis. For example, for transcript-level analyses there are tools to quantify the expression level of transcripts and genes from RNA-seq data [2], identify RNA-binding protein (RBP) binding sites from crosslinking and immunoprecipitation (CLIP) data [3,4], improve transcript annotation with the help of RNA 3’end-sequencing data [5,6], estimate gene expression at the single cell level [7] or improve the annotation of transcripts and quantification of splicing events based on long read sequencing (e.g., on the Oxford Nanopore platform) [8,9]. Such tools are written in different programming languages (e.g., Python, R, C, Rust) and have distinct library requirements and dependencies. In most cases, the tools expect the input to be in one of the widely accepted file formats (e.g., FASTQ [10], BAM [11]), but custom formats are also frequently used. In addition, the variations in protocols or instruments across experiments may make it necessary to use different parameterization for every sample, rendering a joint analysis of samples from multiple studies challenging. Combining tools into an analysis protocol is a time-consuming and error-prone process. As these tasks have become so common, and as the data sets and analyses continue to increase in size and complexity, there is an urgent need for expertly curated, well-tested, maintained and easy-to-use reusable computational workflows.

A number of feature-rich, modern workflow specification languages and corresponding management systems [12,13] like Snakemake [14,15], Nextflow [16] and CWL [17] are now gaining widespread popularity in life sciences, as they make it easier for such workflows to be developed, tested, shared and executed. This leads to more reusable code and reproducible results, while fostering scientific collaborations and Open Source Software along the way. In addition, to facilitate the installation and execution of these workflows across different hardware architectures and host operating systems, modern workflow management systems make use of virtualization and encapsulation techniques relying on containers (e.g., Docker [18] and Singularity [19]) and/or package managers (e.g., Conda [20] and Bioconda [21]). An added advantage of using workflows is the metadata stored along with the expected results. These can be invaluable for re-analyzing the data but may also provide additional insights into the results and cost analyses (e.g., runtimes, resources usage).

The aim of the presented work is the development of a flexible, easy-to use workflow for bulk RNA-seq data processing. The inclusion of the most widely used and best performing tools for the various processing steps minimizes time spent by users on making tool choices. Use of a workflow language for the development ensures the reproducibility and reliable execution of each analysis and it facilitates (meta)data management and reporting.

## Methods/Results

ZARP (**Z**avolan-Lab **A**utomated **R**NA-seq **P**ipeline) is a general purpose RNA-seq analysis workflow that allows users to carry out the most general steps in the analysis of Illumina short-read sequencing libraries with minimum effort. The workflow is developed in Snakemake [14,15], a widely used workflow language [12]. It relies on publicly available bioinformatics tools that follow best practices in the field [22], and handles bulk, stranded RNA-seq data, single or paired-end.

### Workflow inputs

ZARP requires two distinct input files: (1) A tab-delimited file with sample-specific information, such as paths to the sequencing data (FASTQ format), reference genome sequence (FASTA format), transcriptome annotation (GTF format) and additional experiment protocol- and library-preparation specifications like adapter sequences or fragment size. (2) A configuration file in YAML format containing workflow-related parameters, such as results and log directory paths and user-related information. Advanced users can take advantage of ZARP’s flexible design to provide tool-specific configuration parameters via an optional third input file, which allows adjusting the behaviour of the workflow to their specific needs. More information on the input files can be found in ZARP’s documentation [23].

### Analysis steps

A general schema of the workflow in its current version (0.3.0) is presented in Figure 1 (see Supplementary Figure 1 for a more technical representation of the entire workflow, including all of its steps). Table 1 below lists the main tools/functionalities of ZARP:

**Figure 1.**
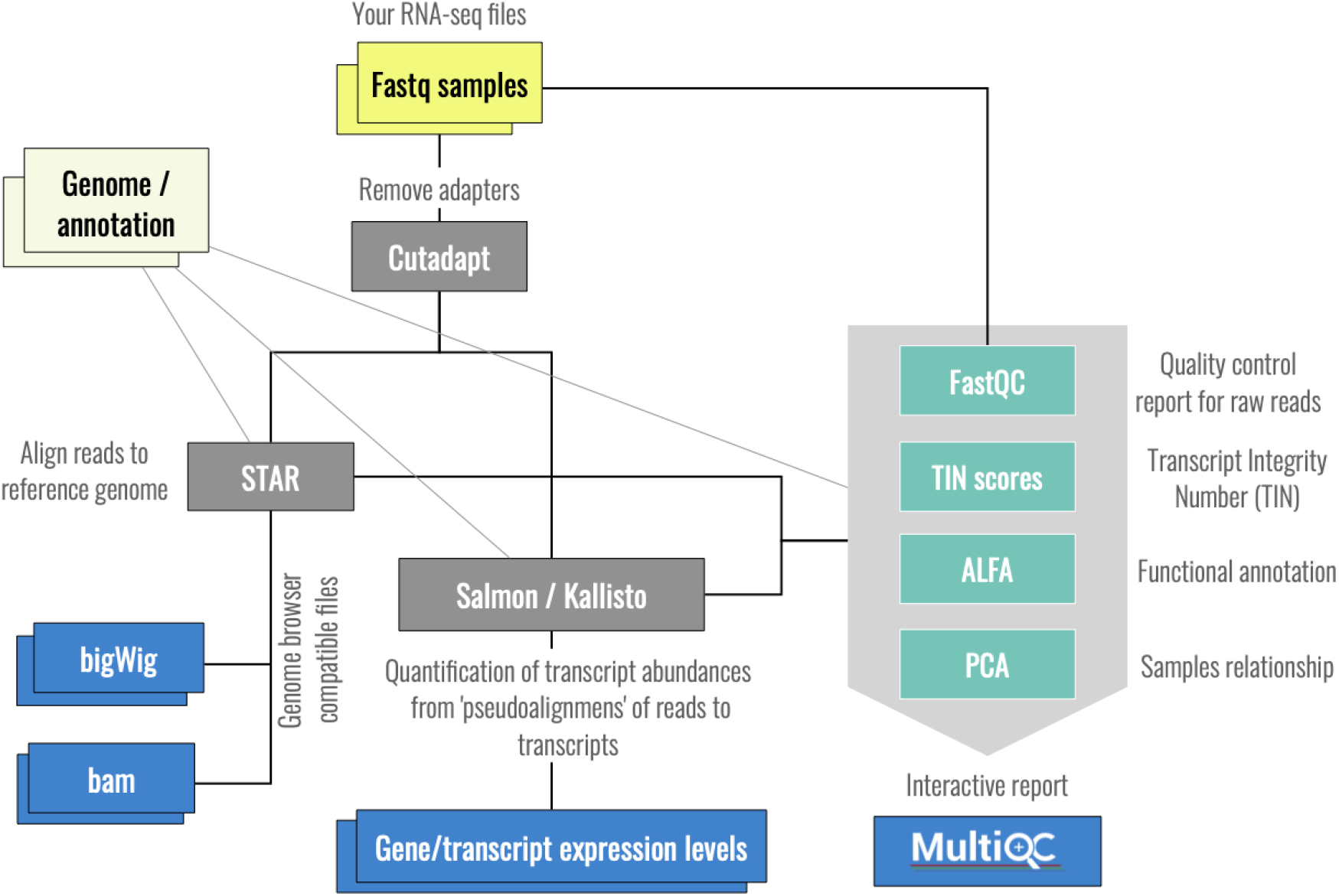
Schematic overview of the ZARP workflow.

**Table 1.** Core tools/functionalities included in ZARP. See main text for more information on use cases for each tool and why we chose those tools to be included in ZARP.

| Tool | Description | Reference |
| --- | --- | --- |
| FastQC | Generates various quality control metrics based on raw FASTQ data. | [24] |
| Cutadapt | Trims sequence fragments of non-biological origin or low information content. | [25] |
| STAR | Aligns reads to reference genome. | [26] |
| tin-score-calculation | Calculates a Transcript INtegrity score (TIN) on aligned reads that reflects the state of RNA degradation of a sample. | [27] |
| ALFA | Annotates read alignments based on gene/transcript annotations. | [28] |
| kallisto | Estimates gene/transcript expression levels. | [29] |
| Salmon | Estimates gene/transcript expression levels. | [30] |
| zpca | Performs principal component analyses of gene/transcript expression level estimates across samples in a given workflow run. | [31] |
| MultiQC | Aggregates tool results and generates interactive reports. | [32] |

Calculation of per-sample quality statistics by applying **FastQC** [24] directly on the input files (FASTQ) provides a quick assessment of the overall quality of the samples. These consist of a considerable range of metrics, including, for example, GC content, overrepresented sequences and adapter content. An excessive bias in GC content may affect downstream analyses and may have to be corrected for [33]. Overrepresented sequences may be the result of PCR duplication, which, if excessive, may skew expression estimates and other downstream analyses. Information about adapter content may be used to cross-check whether it matches with whatever the user has selected to trim. For more information on the metrics that FastQC reports and how they can be interpreted, please refer to [24].

Trimming of any 5’ and/or 3’ adapters as well as poly(A/T) stretches using **Cutadapt** [25] ensures a more reliable alignment as well as removal of contaminant adapter sequences. The adapters and poly(A/T) stretches to be removed are indicated by the user.

Alignment against a given set of genome resources (either only the genome or the genome and a set of corresponding gene annotations) is the step where each read is assigned to the genomic region from which it originated. Even though there are many available aligners, **STAR** has been chosen [26][34]. BAM-formatted files that are then sorted (based on coordinates) and indexed using SAMtools [11]. These steps enable faster random access and visualisation by tools such as genome browsers. The sorted, indexed BAM files are further converted into the BigWig (BedGraphtoBigWig from UCSC tools [35]) format, which allows for library normalisation, and is thus convenient for visualising or comparing coverages across multiple samples.

The aligned reads are also used to calculate per-transcript Transcript Integrity Numbers (TIN scores) [36], a metric to assess the degree of RNA degradation in the sample. This is done with **tin-score-calculation** [27], which is based on a script originally included in the RSeQC package [37] but modified by us to enable multiprocessing for increased performance.

To provide a high-level topographical/functional annotation of which gene segments (e.g., CDS, 3’UTR, intergenic) and biotypes (e.g., protein coding genes, rRNA) are represented by the reads in a given sample, ZARP includes **ALFA** [28].

**Salmon** [30] and **kallisto** [29] along with a transcriptome are used to infer transcript and gene expression estimates. Since both of these tools have been shown to be equally fast, memory efficient and accurate [38], they are both included in ZARP. The main output metrics provided by either tool are estimates of normalized gene/transcript expression, in Transcripts Per Million (TPM) [39], as well as raw read counts per gene/transcript.

Within ZARP, TPM estimates are essential for performing principal component analyses (PCA) [40] with the help of **zpca** [31], a tool created by us for the use in ZARP, but packaged separately so that it can be easily used on its own or as part of other workflows. PCAs on gene/transcript expression levels can help users understand whether differences in gene/transcript expression levels across different sample groups are sufficiently high that meaningful results in downstream analyses may be expected.

TPM and raw count estimates can be further used in downstream analyses, e.g., for differential gene/transcript expression, differential transcript usage or gene set enrichment analyses. Given that such analyses require an experiment design table and are difficult to configure generically for a wide range of experiments, we chose not to include these in ZARP. However, to facilitate downstream analyses, gene/transcript estimates are aggregated for all samples with the aid of Salmon and merge_kallisto [41], which generate summary tables that can be plugged into a variety of available tools.

ZARP produces two user-friendly, web-based, interactive reports: one with a summary of sample-related information generated by **MultiQC** [32], the other with estimates of utilized computational resources generated by Snakemake itself. Note that both fortin-score-calculation and ALFA, we have created plugins so that the respective results can be explored interactively through MultiQC.

### Reproducibility and reusability

To enhance reproducibility of results and reusability of the workflow, each step (referred to as “rule” in Snakemake) of the workflow definition relies either on Conda environments mostly hosted in the Bioconda channel [21] or on Docker images. The latter are converted by Snakemake to Singularity images [19] on the fly where needed, enabling seamless execution of the workflow in environments with limited privileges (e.g., HPC clusters). Users can choose between Conda- and container-based execution by selecting or preparing an appropriate profile when/before running a workflow. At the moment, we include profiles for the Slurm job scheduler and we plan to add new profiles over time. For that, we encourage users to feed their own profiles back to the original ZARP repository so that the entire community can benefit.

### Output and documentation

In addition to the transcript/gene expression tables, ZARP collects log files and metadata for downstream analyses. Intermediate files can be optionally cleaned up by ZARP to minimize disk space usage. The workflow is hosted in its own GitHub repository, and each ZARP version released is accompanied by an up-to-date workflow-oriented description.

### Continuous Integration and Testing

To facilitate collaborative development of the workflow and associated software and to reduce the chance of the codebase regressing with ongoing changes, ZARP is making use of a GitHub Actions-based workflow for Continuous Integration and Delivery (CI/CD). Each modification to the remote repository triggers a variety of integration tests (Conda environments test, Snakemake graph test, dry run, minimal-example based run) to guarantee ZARP’s correct execution throughout the development cycle as the source code is refactored and new features are added.

## Use Cases

Apart from quickly gaining insights into individual samples or smaller sets of samples, ZARP is very well suited to analyze large RNA-Seq experiments or even run meta-analyses across multiple different experiments.

To demonstrate how ZARP can be used to gain meaningful insights into typical RNA-seq experiments, we tested it on an RNA-seq dataset that was generated by Ham et al. (GEO [42] accession number GSE139213) while analyzing the role of mTORC1 signalling in the age-related loss of muscle mass and function in mice [43]. The dataset consists of 20 single-ended RNA-seq libraries (read length: 101 nt, gzipped FASTQ file sizes ranging from 0.8 to 3.2 Gb, library sizes ranging from 18.5 to 75.3 reads), corresponding to four cohorts of 3-months old mice (with five biological replicates per cohort): (1) wild-type, (2) rapamycin-treated, (3) tuberous sclerosis complex 1 (TSC1) knockout and (4) rapamycin-treated TSC1 knockout. The samples were mapped against ENSEMBL’s [44] GRCm38 genome primary assembly and corresponding gene annotations (release: 99) for standard human chromosomes. Other parameters for populating ZARP’s samples table were obtained from the GEO accession entries of the respective samples. Sample tables and results for the test run are publicly available [45].

In Figure 2, we are presenting a subset of the outputs that ZARP generated for this dataset. We can see that the GC content of reads (Figure 2A) is slightly skewed towards being more AU-rich, yet all samples pass the FastQC-defined threshold for GC bias. Moreover, GC content does not exhibit a strong bias across samples. There is no evidence of extensive sequencing of residual adapters (“adapter contamination”) (Figure 2B; black), as less than 1% of reads have been discarded in each sample because of insufficient length after adapter trimming. Transcript integrity across samples is also uniform and high (Figure 2C), with the highest density of expressed transcripts at TIN scores of 75 to 85. Similarly, alignment statistics as reported by STAR are also consistently high (Figure 2D), with rates of reads mapped uniquely against the mouse genome of more than 72% across all samples (<4% unmapped), irrespective of sequencing depth. As expected, ALFA analysis of transcript categories shows that uniquely mapped reads overwhelmingly originated from protein coding genes (over 86% for all samples) (Figure 2E). Taken together, these metrics indicate that all samples are of sufficiently high quality for downstream analyses.

**Figure 2.**
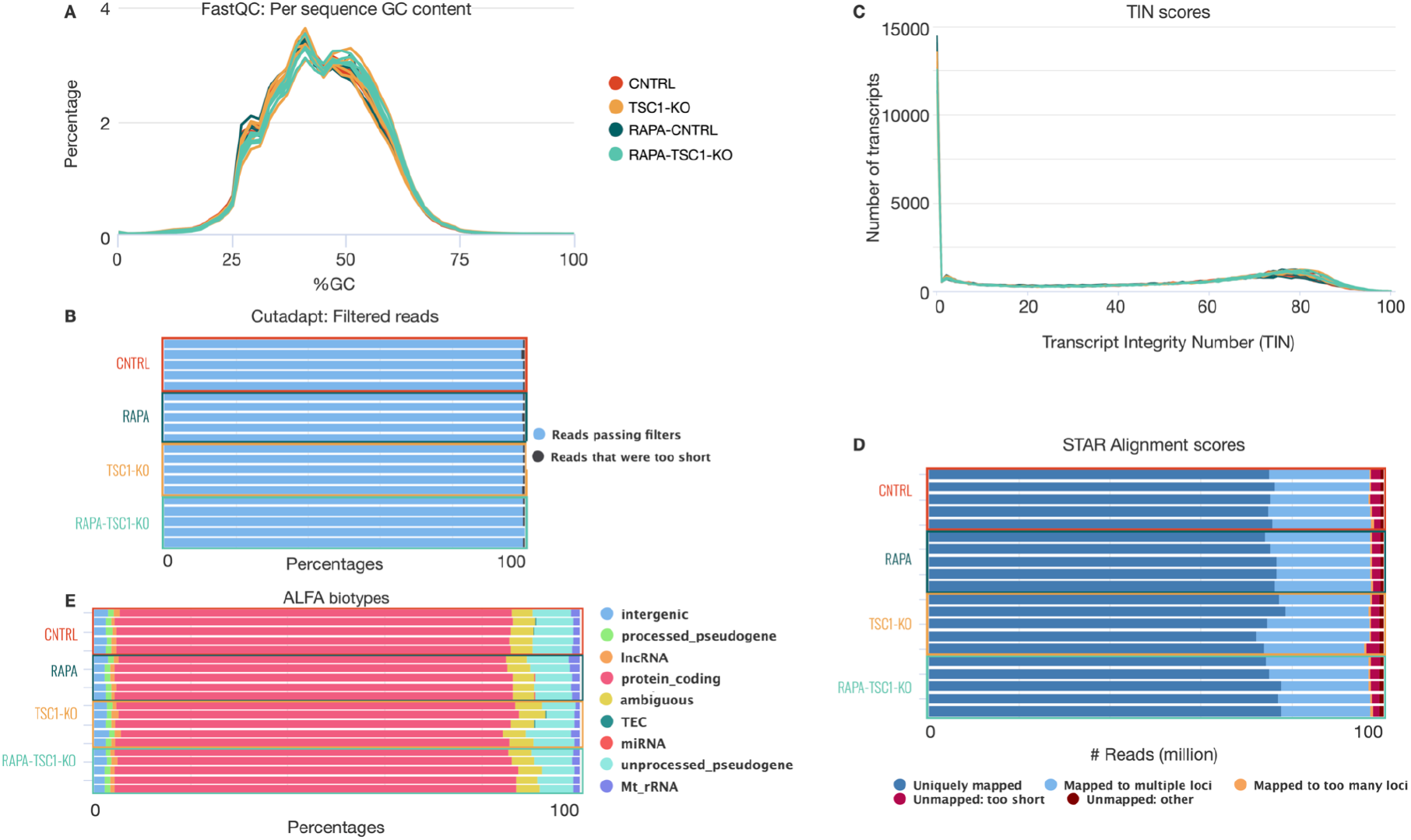
Selection of metrics reported by ZARP. Shown are (A) GC content, (B) adapter removal report, (C) Transcript Integrity Number (TIN) score, (D) STAR alignment statistics, and (E) ALFA biotypes for the test run described in the main text. Figures have been edited for visibility purposes in order to group samples according to cohorts. Additionally, some biotypes have been omitted from (E) as they are not meaningfully represented. Note that in (C), transcripts that are not expressed are assigned a TIN score of 0. The complete raw html report can be found at [45].

In addition to sample-specific metrics, ZARP also provides tooling to compute principal component analyses across samples (Figure 3). For the test run, the distribution of samples in the space of the first two principal components shows a clustering by condition, with a clear separation between knockout and wild type, as well as between the untreated and rapamycin-treated TSC1 knockout mice. This separation is more pronounced at the gene expression level (Figure 3A), but is also present at the transcript level (Figure 3B). This shows that the differences across conditions are more pronounced than any replicate biases (multiplicative noise, sequencing errors), i.e., the signal-to-noise ratio is favorable, which strongly increases the likelihood that any subsequent analyses (e.g., differential gene/transcript expression analysis) will provide targets of biological importance.

**Figure 3.**
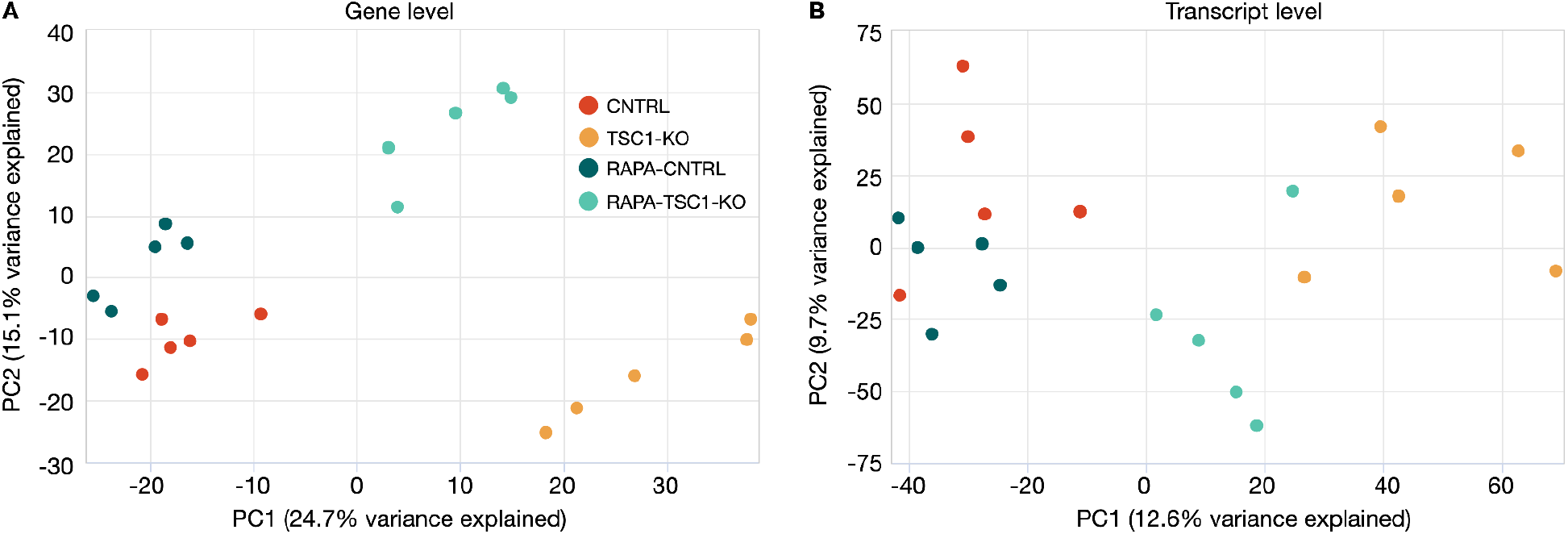
Principal component analysis. Principal component analysis (PCA) at the (A) gene and (B) transcript level. PC1 and PC2 correspond to the first and second principal components, respectively. Variances explained by each of them are stated in the parentheses of the corresponding axes labels. Expression levels used in this figure are those reported by kallisto, but ZARP also generates corresponding PCA plots for Salmon-based quantifications.

The total wall clock time to execute the entire test run was just over one hour (1.01h) for all 20 samples on our Slurm-managed HPC cluster [46], where we could make heavy use of ZARP’s parallelization capabilities. This translates to a total CPU time of 68.79 h, out of which 6.68h were run-specific, i.e., jobs that had to be executed only once for all samples. The accumulated sample-specific CPU time used for each sample varied between 2.75h and 8.44h. While the actual runtime may differ considerably across different compute environments, we project that most users would be able to run even large-scale analyses with dozens to hundreds of samples in less than a day on an HPC cluster, with very little hands-on time. Maximum memory usage for any of the steps and across all samples was <32 Gb (for STAR indexing and mapping of/against the human genome), indicating that ZARP is suitable for execution on state of the art computers, albeit at considerably higher runtimes due to limited parallelization capabilities, particularly for large sample groups. None of the jobs took longer than ~20 min (wall clock time) for any of the samples (Figure 4). Among the most time-consuming steps are the creation of indices (STAR, Salmon, kallisto), which however have to be performed only once per set of genome resources. Among the sample-specific steps, the calculation of the Transcript INtegrity (TIN) score was the most time-consuming. However, we had already considerably reduced its runtime by adding parallelization capabilities to the original script (see subsection “Analysis steps” in section “Methods/Results” for details).

**Figure 4.**
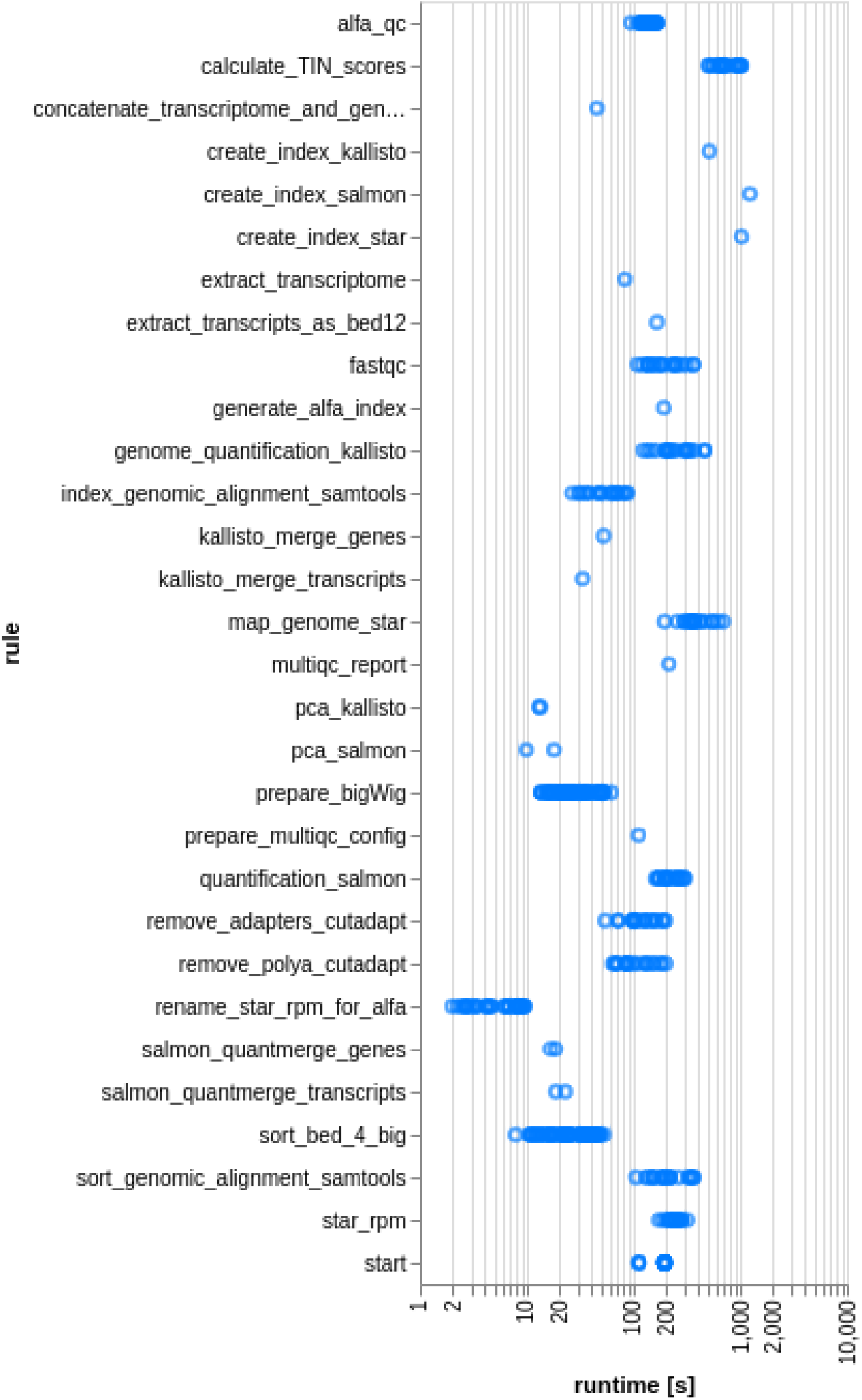
Runtime statistics. Runtime (in seconds; wall clock time) of the different steps (“rules”) of the workflow run are depicted for each sample. The workflow was executed in an HPC cluster managed by the Slurm job scheduler, so the reported runtimes include the time that jobs spent queuing. Additional variation in runtimes may result from individual jobs being executed on cluster nodes with different specifications.

In summary, our test case demonstrates how ZARP can be used to quickly gain informative insights (Figures 2 & 3) into a non-trivial real-world RNA-seq analysis in a reasonable timeframe (Figure 4).

## Discussion/Conclusions

ZARP is a general purpose, easy-to-use, reliable and efficient RNA-seq processing workflow that can be used by molecular biologists with minimal programming experience. Scientists with access to a UNIX-based computer (ideally a Linux machine with enough memory to align sequencing reads) or a computing cluster can run the workflow to get an initial view of their data on a relatively short time scale. ZARP has been specifically fine-tuned to process bulk RNA-seq datasets, allowing users to run it out of the box with default parameters. At the same time, ZARP allows advanced users to customize workflow behavior, thereby making it a helpful and flexible tool for edge cases, where a more generic analysis with default settings is unsuitable. The outputs that ZARP provides can serve as entry points for other project-specific analyses, such as differential gene and transcript expression analyses. ZARP is publicly available and open source (Apache License, Version 2.0), and contributions from the bioinformatics community are welcome. Please address all development-related inquiries as issues at the official GitHub repository [47].

## Data and Software Availability

### Data

Raw data analysed in section “Use Cases” are publicly available for anyone to download from the NCBI:GEO server, accession number GSE139213.

### Software

The ZARP code is available on GitHub at [47] and is published under Apache License, Version 2.0. A snapshot of the ZARP version described in this manuscript (0.3.0) has been additionally uploaded to Zenodo for long-term storage [23]. Both services are public and allow anyone to download the software without prior registration.

### Results

Analysis results presented in section “Use Cases” are publicly available for anyone to download from Zenodo.

## Author Contributions

MK, FG, MZ, AK conceived the project. MK, FG, CJH, DB, MB, PI, AK developed the method. MK, FG, CJH, DB, MB, PI, KA, MZ, AK developed custom tools used in the study. MA, AB tested the method with real datasets. MK, FG, CJH, DB, MB, PI, MA, MZ, AK wrote the manuscript. MK, MZ, AK supervised the study. MK, MB, AK managed the software repository. All of the authors approved the manuscript.

## Competing Interests

None declared.

## Grant Information

M.K., M.B., D.B. were supported by Biozentrum Basel International Ph.D. Program Fellowships for Excellence.

## Acknowledgements

This work would not have been possible without the sciCORE Team [46] at the University of Basel. We are thankful for their support regarding the computational infrastructure as well as dedicated time and effort to aid us in this project. We would also like to express our deepest gratitude towards all members of the Zavolan Lab who contributed to this work with numerous pieces of advice, during the initial development as well as by testing the workflow in later stages. We would also like to thank the bioconda community for helping us package and distribute some of the custom tools we developed.

## Supplementary material

**Supplementary Figure 1.**
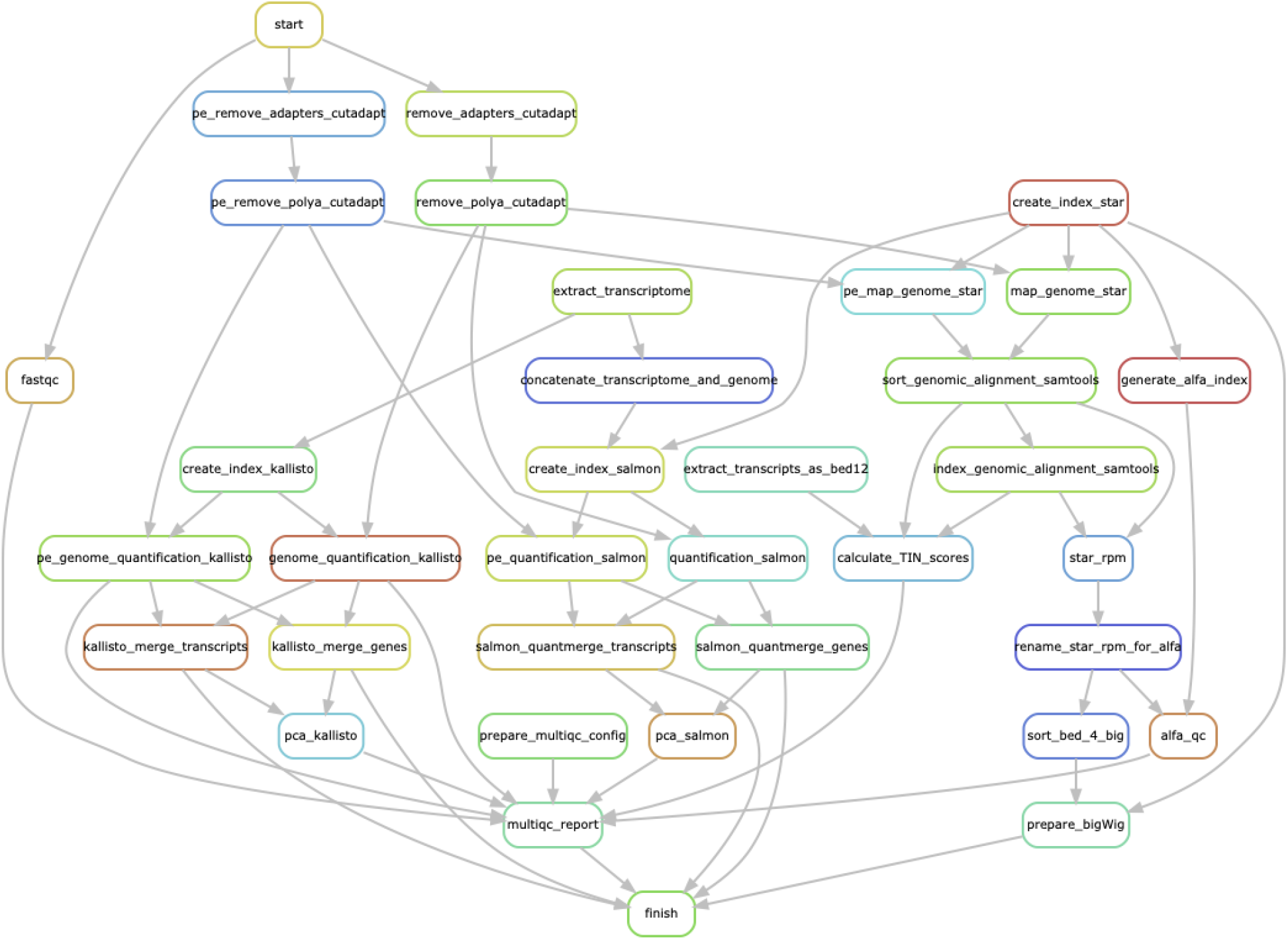
ZARP workflow schema. Graph-based representation of ZARP v0.3.0, including all of its steps (“rules”), as produced by running Snakemake with the `--rulegraph` option. Steps for both the single and the paired end workflows are shown.

## Notes

https://github.com/zavolanlab/zarp

https://zenodo.org/record/5703358

https://zenodo.org/record/5683524

### Competing Interest Statement

The authors have declared no competing interest.

